# Common drugs alter microbial protein expression, but not composition in fecal cultures from Crohn’s disease patients

**DOI:** 10.1101/2023.08.19.553599

**Authors:** Heike E.F. Becker, Ronny Mohren, N. Giang Le, Luc J.J. Derijks, Daisy M.A.E. Jonkers, John Penders

**Author notes:** Corresponding authors: Daisy Jonkers, John Penders. authors contributed equally (Daisy M.A.E. Jonkers, John Penders).

## Abstract

**Introduction:** A substantial number of Crohn’s disease (CD) patients experience side-effects and/or non-response to medical drugs. In part, this might be attributed to the interaction of the intestinal microbiome with xenobiotics. The aim of this study was to explore the effect of common CD drugs on the patient’s microbiome *in vitro*.

**Methods:** The fecal microbiome of each of 5 CD patients was exposed to 42 μg/ml budesonide, 55 μg/ml 6-mercaptopurine (6-MP), 5 or 15 μg/ml tofacitinib, or DMSO-control in defined culture medium in an anaerobic chamber at 37°C for 24 hours. Subsequently, DNA and proteins were isolated and subjected to 16 rRNA gene amplicon sequencing and LC-MS proteomic analysis, respectively.

**Results:** Metagenomic and metaproteomic analyses revealed larger differences between donors than between drug exposures. Exposure to 6-MP and tofacitinib resulted in a significant alteration in the metaproteome when compared to the control condition, whereas no effect could be observed for budesonide. Applying a stringent selection, 33 proteins were more abundant and 93 less abundant in all 6-MP cultures and could thereby discriminate clearly between 6-MP and control. In contrast to metaproteomic analyses, metagenomic analyses only detected a lower relative abundance of *Colidextribacter* in 15 μg/ml tofacitinib cultures, but not in overall richness, diversity or community structure.

**Conclusion:** Tofacitinib and especially 6-MP clearly affect microbial function, but barely microbial composition *in vitro*. These drug-induced functional changes may subsequently influence host physiology and potentially inflammation in CD. Our findings emphasize the relevance to include functional microbial studies when investigating drug-microbiota interactions. Further research needs to elucidate the impact of 6-MP-induced microbial alterations on intestinal physiology and inflammation in CD.

## Introduction

Crohn’s disease (CD) is a chronic inflammatory gastrointestinal disease with an alternating disease course^1,2^, characterized by flares and periods of remission. There is no cure for CD and medical treatment is focused on the induction and subsequent maintenance of remission. Recommended drugs include local and systemic corticosteroids, thiopurines, and anti-TNF biologicals^3^.

Although the pathophysiology of CD is not completely elucidated, several factors are associated with disease onset. These include a genetic predisposition, environmental factors, an aberrant immune response to the commensal microbiota, intestinal barrier disruption, and microbial dysbiosis. The past years, the latter gains increasing attention and has shown to be related to disease onset as well as disease course^4–6^. In the healthy situation, the intestinal microbiota contributes to immune development and function, an intact mucosal barrier, and digestion of food. Although the intestinal microbiota composition varies widely between individuals, the microbiota of CD patients is characterized by a decreased fecal and mucosal diversity, decreased temporal stability, and a shift in the abundance of specific bacterial taxa^7^. Alterations in microbial function might, however, have an even larger impact on host health and intestinal inflammation than alterations in microbiota composition^8^.

Recent studies have demonstrated that common medical drugs can impact intestinal microbial composition and growth. For instance, Maier *et al*. identified 203 human targeted drugs, which affected the growth of 40 isolated bacterial strains^9^. Thereby, drug-induced alterations in microbiota composition and function can potentially contribute to mucosal healing, drug response, or drug-related side effects^10^.

Also, for the various drug classes used in the treatment of CD, changes in microbiota composition or bacterial growth have been detected, although the specific effects of these drugs on the microbiota and related clinical effects are difficult to interpret due to the heterogeneity in research methodology^11^. For instance, several studies investigated the thiopurines azathioprine and 6-mercaptopurine (6-MP) on intestinal microbes using different study designs. While an effect on bacterial growth or abundance has been found consistently, the exact outcomes differed. Together, 6-MP is found to inhibit bacterial growth of, for instance, *Prevotella copri, Bacteroides fragilis* and *Eggerthella lenta*^9,12,13^. Some of the studies also reported drug-microbiota interaction with other drugs that are relevant in CD treatment, including budesonide and prednisolone^14–17^.

The vast majority of studies have investigated the microbiota composition, whereas the impact of drugs on microbiota function remains largely unexplored. However, compositional changes may not predict the nature of the subsequent functional changes, which can finally influence, for instance, intestinal inflammation, but may also impact microbial drug metabolism and thereby drug response and toxicity. Therefore, we aim to explore the impact of the frequently used drugs budesonide, 6-mercaptopurine, and tofacitinib on CD patient microbiota function in an *ex vivo* fecal culture study and to compare the extent of functional variation with the compositional changes. We chose to select these drugs, because they cover three different drug classes and working mechanisms. In addition, budesonide and 6-MP are frequently used in CD treatment, and tofacitinib is a novel drug, shown to be effective in CD treatment^3,18^.

## Methods

### Study population

Five CD patients from the IBD South Limburg (IBD-SL) cohort kindly provided a fresh fecal sample and completed a short questionnaire on demographics and potential confounders, including medication use and body weight. Prior to study participation, all participants gave written informed consent. The IBD-SL cohort was approved by the Medical Ethics Committee of the Maastricht University Medical Centre+ (IBD-SL: NL31636.068.10) and registered on www.clinicaltrials.gov (NCT02130349). The study was conducted in accordance with the revised Declaration of Helsinki and in compliance with good clinical practice.

### Sample collection and fecal culture

The participants collected complete defecations, using airtight boxes and one sachet AnaeroGen 1.5 L (Ref.: AN0025A, ThermoScientific). The sample was stored immediately at 4 °C until processing within 5 hours. The sample was transferred into an anaerobic chamber with 80 % Nitrogen, 10 % CO_2_, and 10 % H_2_ and homogenized in defined culture medium as reported by Li *et al*. with pH 7.8^13^. The final incubations contained 2 % (w/v) fecal sample, 1 % dimethylsulfoxide, and were exposed to either 55 μg/ml budesonide (Ref.: B7777, Sigma-Aldrich), 42 μg/ml 6-mercaptopurine (Ref.: 852678, Sigma-Aldrich), 5 μg/ml (10 %) or 15 μg/ml (30 %) tofacitinib (Ref.: PZ0017, Pfizer®) or medium. We chose these concentrations based on the maximum oral intake distributed in 200 g colon content. Since 6-MP is dosed per kg body weight, we based our calculations on the approximate median body weight of CD patients, namely 70 kg^19^. All conditions were incubated in duplicate in a 2 ml 96-deep well plate covered with a PCR cover slip with ventilation holes and being placed in an IKA MS 3 basic plate shaker at 500 rpm for 24 hours at 37°C in the anaerobic chamber. Thereafter, samples were processed on ice and large debris removed by three times repeated centrifugation at 300 x g for 5 min. Then, the culture was pelleted and washed with PBS at 2272 x g for 1 hour at 4 °C as described by Li *et al*^13^. DNA was isolated as described elsewhere^20^, following the adapted protocol Q as described by the Human Microbiome Standards consortium^21^. Proteins were isolated by diluting washed pellets in 1 ml 0.01 % RapiGest SF (Ref.: 186001861, Waters) in 50 mM ammonium bicarbonate as done for DNA isolation. The supernatant was harvested after mechanical lysis and centrifugation at 16000 *g*.

### Microbiota profiling

To identify alterations in microbial composition due to 24-hour drug exposure, we profiled the fecal microbiota at t = 24 hours using 16S rRNA gene amplicon sequencing. The procedure has been described in our previous work by Galazzo *et al*^22^. In brief, the hypervariable V4 region was PCR amplified using the primer pair 16S-341_F and 16S-805_R (CCTACGGGNGGCWGCAG and GACTACH VGGGTATCTAATCC, respectively). After clean-up using AMPure XP purification (Agencourt, MA, USA), the amplicons were quantified using Quant-iT PicoGreen dsDNA reagent kit (Thermo Fisher Scientific, Landsmeer, the Netherlands) and pooled at equimolar concentrations. Finally, amplicons were sequenced on an Illumina MiSeq instrument.

Pre-processing of sequencing data was performed using an in-house pipeline based upon DADA2^23^ that consisted of the following steps: reads filtering, identification of sequencing errors, dereplication, inference and removal of chimeric sequences. In order to assign taxonomy, DADA2 was used to annotate down to the species level using the SILVA 138 version 2 database^24^. Decontam was used with the “either” setting, combining prevalence and frequency-based identification of contaminating Amplicon Sequence Variants (ASVs)^25^. Contaminating sequences were filtered out as well as ASVs present in less than 20 % of all samples and ASVs with an overall abundance below 0.01 % across samples. Data were further analyzed using MicrobiomeAnalyst^26^ for beta diversity indices and GraphPad Prism version 5 for alpha diversity indices. Statistical analyses are stated in the respective paragraphs. Differences in relative abundances were analyzed by LinDA ^27^ version 0.1.0 using R version 4.1.2^28^.

### Proteomics analysis

To explore microbial proteins and enzymatic pathways linked to drug exposure, we conducted proteomics analysis based on liquid chromatography followed by label-free quantitative mass spectrometry (LC-MS).

Isolated proteins were reduced with 20 mM Dithiothreitol (DTT) for 45 minutes and alkylated with 40 mM Iodoacetamide for 45 minutes in the dark. Alkylation was terminated by 20 mM DTT. Subsequent protein digestion was performed using Endoproteinase LysC and Trypsin (Ref.: V5071, Promega), diluted 1:25 (enzyme to protein) for 2 hours at 37 °C and 750 rpm. For further overnight digestion, 50 mM ammoniumbicarbonate (ABC) was added to reach a final concentration of 1 M urea, and terminated by 1 % formic acid. LC-MS analysis was performed as described by Mezger *et al*.^29^, adapted slightly using a 110 minute gradient from 4 % to 32 % acetonitrile.

For protein annotation, the human microbiome database was used as described by Li *et al*.^30^ and analyzed using Proteome Discoverer 2.5 (Thermofisher). For subsequent data analysis, False Discovery Rate (FDR) corrected (FDR < 1 %) and normalized values were used. KEGG orthology was applied for functional annotation using GhostKOALA with option “prokaryotes”^31^. Microbial taxonomic annotation based on identified peptides was conducted using Unipept 4.0^32^.

## Results

### Patient population

The baseline characteristics of the CD feces donors are described in table 1. One patient used the thiopurine pro-drug azathioprine at the time of donation, which is usually metabolized to 6-thioguanine nucleotides via 6-MP in the liver^33^. The other patients did not use any of the investigated drugs, *i*.*e*. budesonide, 6-MP, or tofacitinib.

**Table 1:** Baseline characteristics of study participants.

|  | CD1 | CD2 | CD3 | CD4 | CD5 |
| --- | --- | --- | --- | --- | --- |
| Sex (F/M) | Male | Female | Male | Male | Female |
| Age in years | 27 | 45 | 38 | 43 | 41 |
| BMI kg/m<sup>2</sup> | 21.0 | 42.4 | 31.4 | 23.1 | 23.4 |
| HBI | 1 | 0 | 0 | 1 | 1 |
| BSS | 3 | 4 | 5 | 1 | 5 |
| IBD medication | Vedolizumab | None | Vedolizumab | Adalimumab | Azathioprin,<br>Mesalazin |
| Antibiotic use in past 3 months (n) | No | No | No | No | No |
| PPI use (n) | No | No | No | No | No |
| Medication change in last week (n) | No | No | No | No | No |
| Probiotic use in last week (n) | No | No | No | No | No |
| Alternative diet (n) | No | No | No | No | No |
| Current smoker (n) | No | No | Yes | No | No |
| Current pregnancy (n) | No | No | No | No | No |

BMI = body mass index; HBI = Harvey Bredshaw Index; BSS = Bristol Stool Scale PPI = proton pump inhibitor

**Table 1:** Comparison of microbial richness and diversity between the experimental conditions.

| Comparison | Observed richness p-value | Shannon index p-value |
| --- | --- | --- |
| Ctrl vs Budesonide | 0.586 | 1.000 |
| Ctrl vs 6-MP | 0.345 | 0.125 |
| Ctrl vs Tofacitinib 10 % | 0.188 | 0.625 |
| Ctrl vs Tofacitinib 30 % | 0.136 | 0.313 |

### The impact of 24-hour fecal culture on microbiota composition

To evaluate the effect of the 24-hour fecal culture system in itself, we compared the microbiota composition of the original fecal samples with the respective 24-hour cultures using 16S rRNA gene amplicon sequencing. The median difference in observed richness between the original samples was 8 ASVs (range -80; 92). The respective differences in bacterial diversity analyzed by the Shannon index were -0.373 (range -0.174; -0.415) versus 0.241 (range -0.841; 1.163). After 24 hours of culturing, the observed bacterial richness increased with a median of 3 taxa (range: -8; 13) as compared to the respective original fecal sample. Thus, the difference in richness and diversity between original samples was larger as compared to the effect of the 24-hour culture. Using the Wilcoxon signed rank test in GraphPad Prism v5, the observed richness (Figure 1A) and Shannon index (Figure 1B) did not differ significantly between the original fecal samples and 24-hour culturing (p = 0.625; p = 0.063, respectively). The Bray-Curtis index, comparing the microbial community structure, was also not significantly different between original fecal samples and 24-hour cultures (p for PERMANOVA = 0.152; Figure 1C). Together, the effect of the culture was unlikely to bias the experimental outcomes and conclusions. For all downstream analyses, we only analyzed the 24-hour cultures.

**Figure 1:**
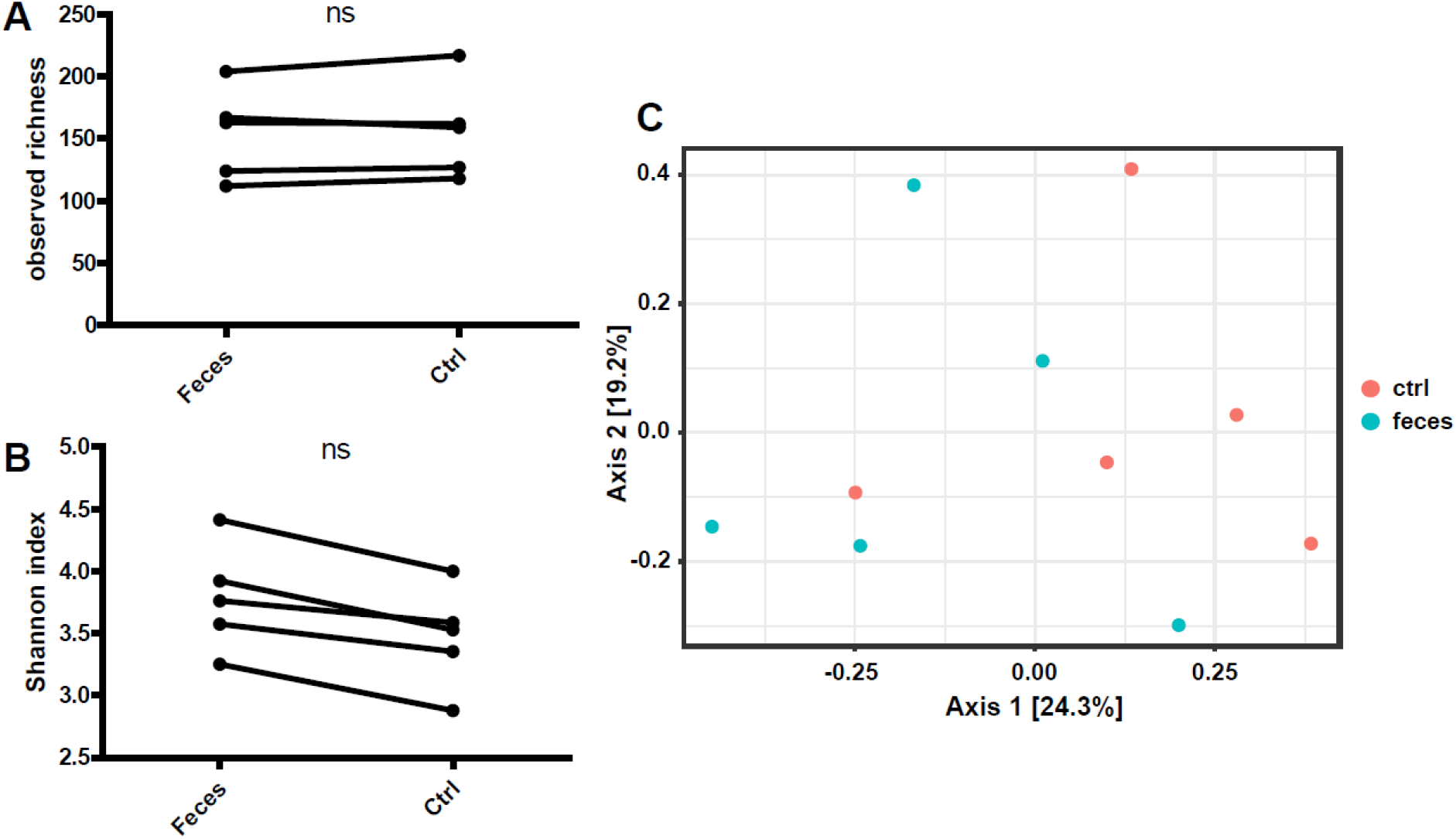
The effect of ex vivo culture on microbiota diversity. The observed richness (A) and the Shannon index (B) were not significantly different before and after 24-hour culturing (p = 0.625; p = 0.063, respectively). The Bray-Curtis index was not significantly different between the original fecal samples and the respective 24-hour cultures (p = 0.152).

### The impact of 24-hour drug exposure on microbiota composition

To detect differences in microbiota composition as a response to different IBD drugs, we profiled the fecal cultures derived from the five CD patients after 24 hours. The observed richness (Figure 2A) and Shannon index (Figure 2B) were not significantly different when comparing the control condition with each experimental condition, using Wilcoxon signed rank test in GraphPad v5 (Table 1). When comparing the donors, the observed richness (Figure 2C) and Shannon index (Figure 2D) were significantly different between donors (both p < 0.001) using Kruskall-Wallis test. Also the Bray-Curtis index was significant when comparing donors (p = 0.001; Figure 2E), but not when comparing the experimental conditions (p = 1, Figure 2E). Further, comparing the relative abundances of bacterial genera between the control condition and each experimental condition, *Colidextribacter* was significantly decreased in the 24-hour cultures exposed to tofacitinib 30 % (p < 0.05) and *Eggerthella* was decreased in the cultures with 6-MP (p < 0.1; Figure 2F).

**Figure 2:**
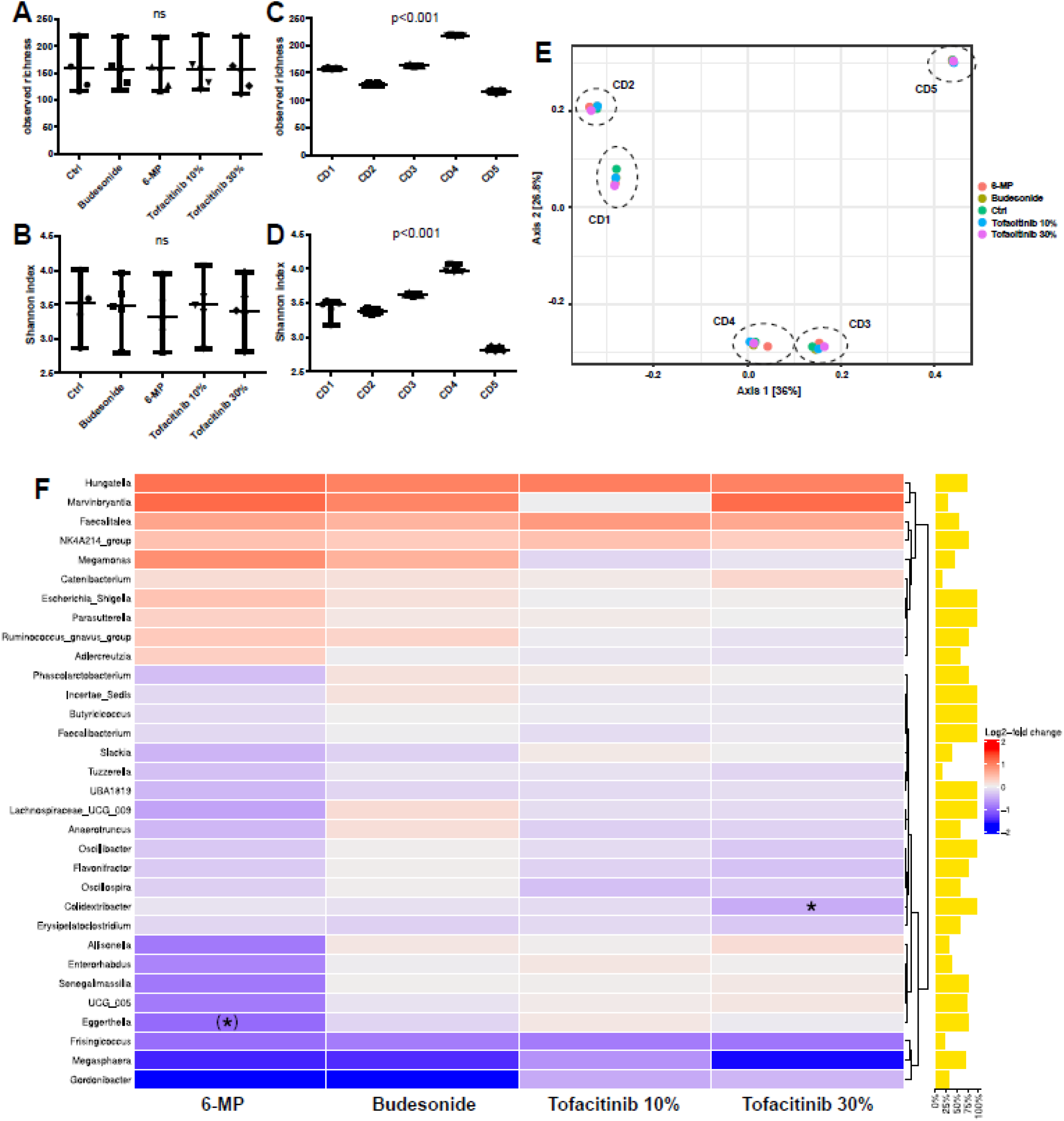
Alterations in microbial composition upon exposure to budesonide, 6-mercaptopurine, or tofacitinib. Observed richness (A) and Shannon index (B) between experimental conditions (none significant, see table 1) and between donors (both p < 0.001; C, D, respectively). Principle Coordinate Analyses on Bray-Curtis dissimilarity (E). Experimental conditions, indicated by the different colors, did not result into significant clustering of samples (p = 1.0), whereas samples from the same donor, highlighted by the dashed ellipses, did significantly cluster together (p = 0.001). Heatmap of log2-fold changes in relative abundances of bacterial genera upon drug exposure as compared to the control condition (F). The prevalence of the genera is depicted as vertical bars. * = p < 0.05; (*) = p < 0.1

### Proteomic profiling of drug exposure on fecal cultures from one CD patient

To explore the impact of budesonide, 6-MP, and tofacitinib (30 %) on microbial protein expression, we performed microbial proteomics analysis with all experimental conditions of one CD patient in duplicate. The patient was chosen randomly. Principal component analysis showed a clear distinction between 6-MP and control, and between tofacitinib and control, whereas budesonide could not be discriminated from the control condition (Fig. 3A). Hierarchical clustering analysis identified 6-MP as being most distinct from the control conditions (Fig. 3B). Therefore, we decided to further explore the impact of 6-MP on microbial protein expression cultures for all five CD patients.

**Figure 3:**
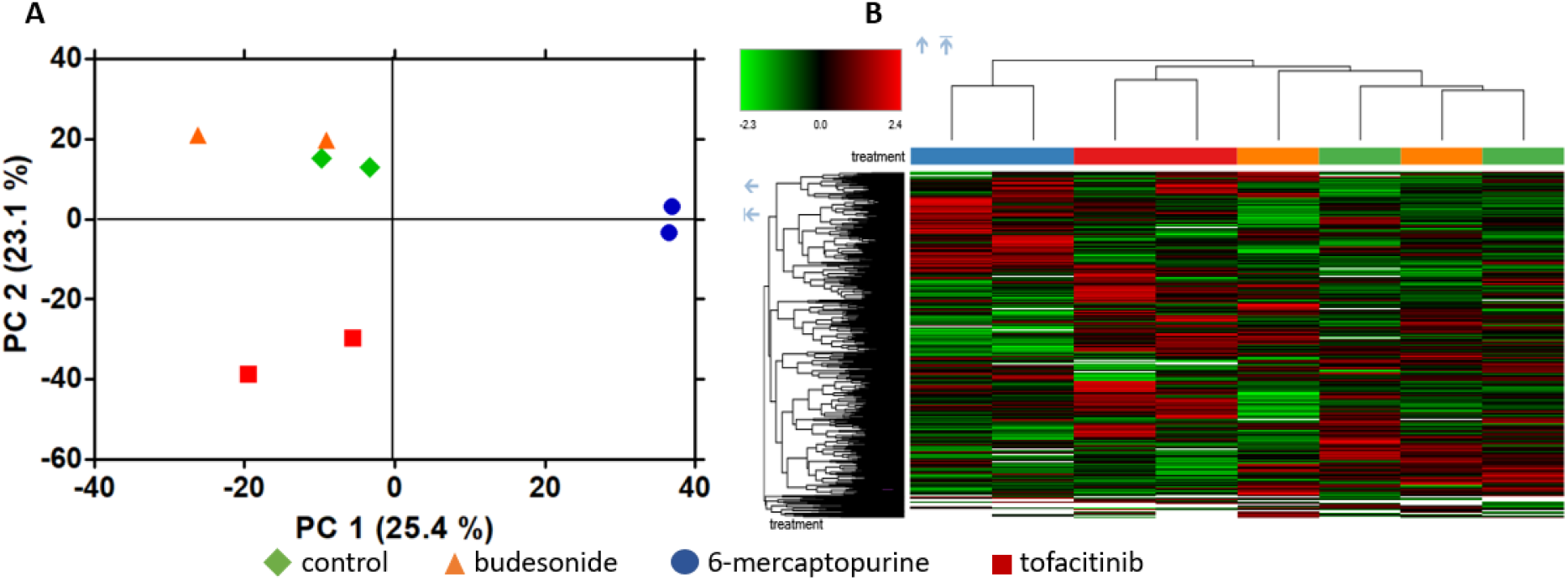
Proteome-based differences between drugs within fecal cultures of one CD patient. Principal component analysis (A) and heatmap (B) comparing the impact of budesonide, 6-mercaptopurine, tofacitinib, and control on the fecal microbial protein expression of one CD patient after 24 hours incubation in duplicate.

### Proteomic profiling of 6-MP exposure on fecal cultures from five CD patients

Principal component analysis revealed a clear distinction based on 6-MP exposure by principal component #5 (Fig. 4A), while the origin of the microbial cultures again had the largest impact on alterations of the metaproteome (Fig 4B). Hierarchical clustering analysis confirmed a higher discrimination based on the donating patients, though with consistent discrimination based on 6-MP exposure for each patient’s fecal culture (Fig 4C).

**Figure 4:**
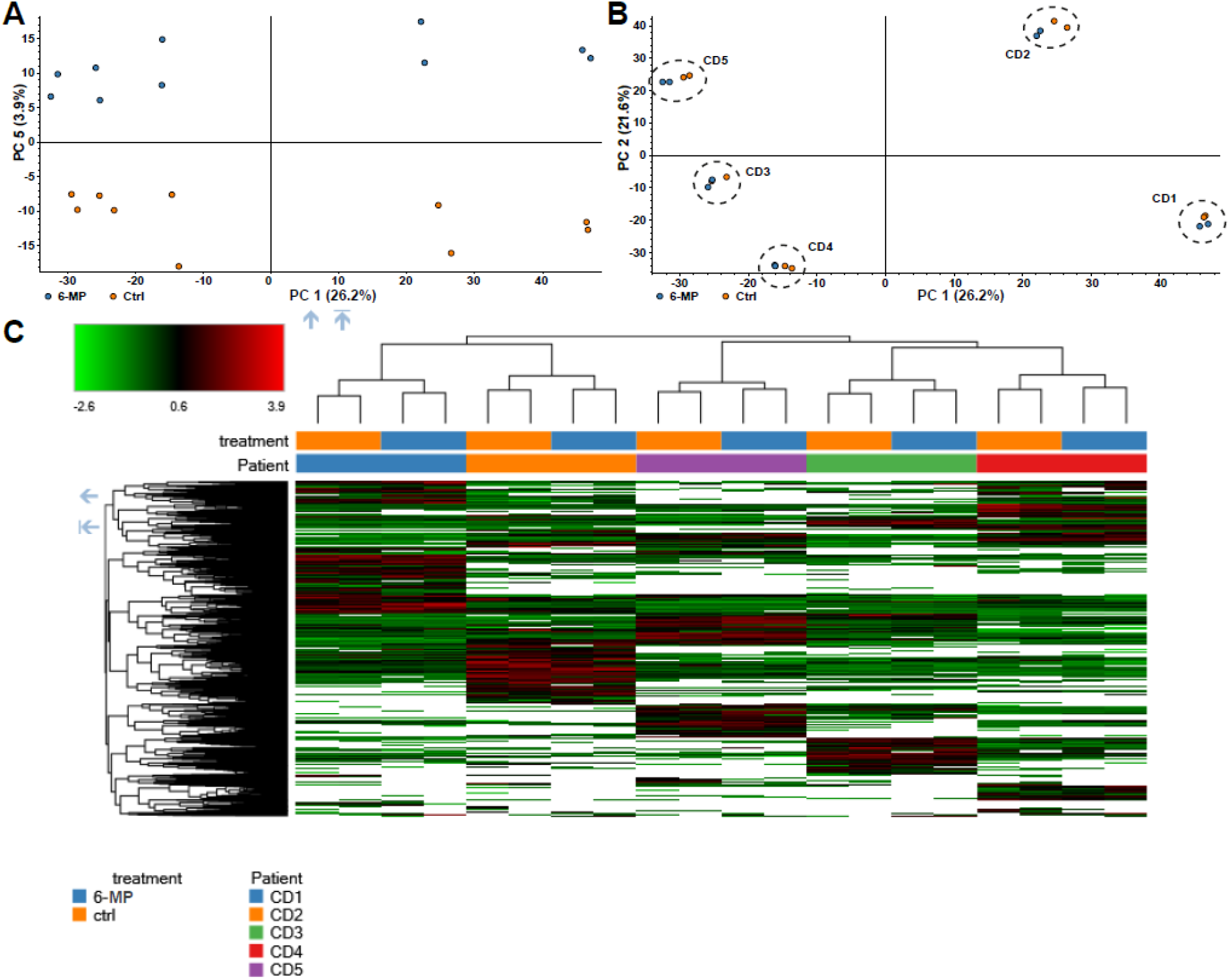
Metaproteome-based differences between 6-mercaptopurine exposure and control comparing five CD patients. Principal component analysis discrimination based on 6-MP exposure in PC 1 and PC 5 (A), and based on donor in PC 1 and PC 2 (B). Heatmap of protein abundances (C) comparing the effect of 6-MP and control on the fecal cultures.

**Figure 5:**
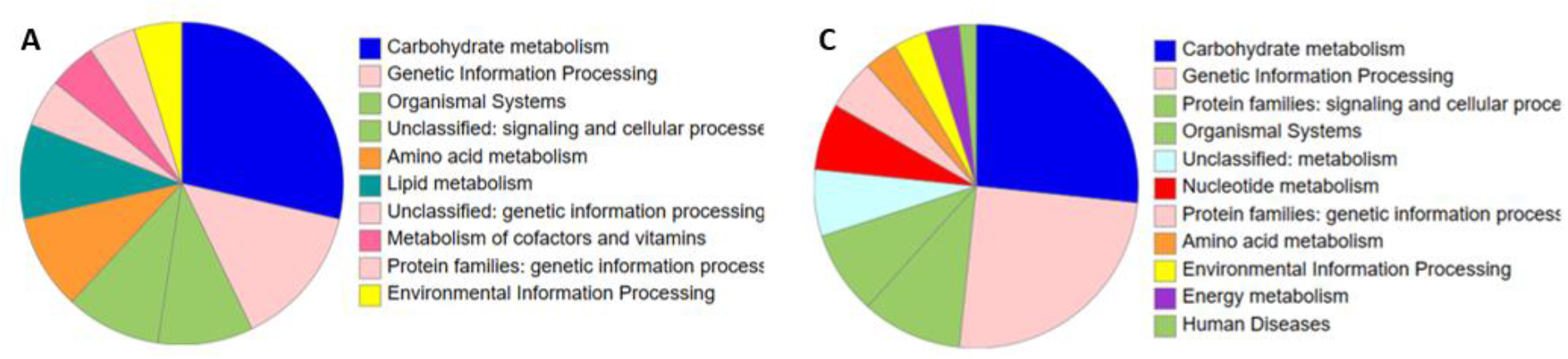
Functional annotation of 6-MP associated proteins. Pie charts displaying microbial proteins that were increased (A) or decreased (B) due to 6-MP exposure. Proteins were annotated using KEGG orthology assignment with the GhostKOALA^31^ online tool.

To further investigate altered biological processes and to identify bacterial proteins that are increased or decreased due to 6-MP exposure, we performed differential expression analyses of protein abundance. We applied a stringent filtering to limit the chance of spurious findings. To this end, we only included proteins present in all donors and in at least 80% of the 6-MP exposed samples when examining the enrichment of proteins upon drug exposure. When examining the depletion of proteins upon drug exposure, we only included proteins that were present in all donors and at least in 80% of the control samples.

After 6-MP exposure, 33 proteins were significantly increased (> 1.5-fold change) and 93 proteins were significantly decreased of the 4178 proteins that were identified by Proteome Discover from all samples in total. Applying KEGG orthology assignments using the GhostKOALA^31^ online annotation tool, 63.6 % of the increased proteins could be annotated (Fig 4A, Supplementary table 1). In general, several proteins were annotated to different pathways. Most of these were found in pathways related to carbohydrate metabolism (six proteins), covering different physiological processes, including glycolysis, citrate cycle, and pyruvate metabolism. Second most common proteins were annotated to the ribosome (three proteins; Fig 4B).

Of the decreased proteins, 64.5 % could be annotated. Most of the decreased proteins were also found to be part of carbohydrate metabolism (16 proteins), mainly including glycolysis, citrate cycle, and pyruvate metabolism (Fig 4C, Supplementary table 2). Second most decreased proteins were related to genetic information processing (15 proteins), most of them to the ribosome (eight proteins; Fig 4D). Besides the ribosome, seven proteins were annotated to amino acid metabolism, two to translation factors, and one as aminoacyl-tRNA biosynthesis, which together are all involved in protein synthesis. Further, three proteins were related to purine metabolism and two to pyrimidine metabolism, which is both related to DNA synthesis and bacterial growth. Moreover, three proteins are related to transporters, of which one was annotated as trimeric autotransporter adhesin, which is related in bacterial adhesion^34^.

### Microbial taxonomic annotation of 6-MP associated proteins

The proteins that were found to be increased or decreased in at least 80 % of the samples due to 6-MP exposure, could be related to specific microbial taxa using Unipept 4.0^32^. For this analysis, the predicted proteins by Proteome Discoverer were used, as submitting the original peptide sequences yielded only very low annotation rates. Taxa associated with the proteins that were increased in all donors due to 6-MP exposure mainly belonged to the phyla Bacteroidetes, mainly *Prevotella*, Firmicutes and Proteobacteria, mainly Sutterellaceae, and were present in comparable abundances (Fig. 6A). In contrast, proteins that were decreased, mainly belonged to Bacteroidetes, mainly *Bacteroides*, and in lower numbers to Firmicutes (Fig. 6B).

**Figure 6:**
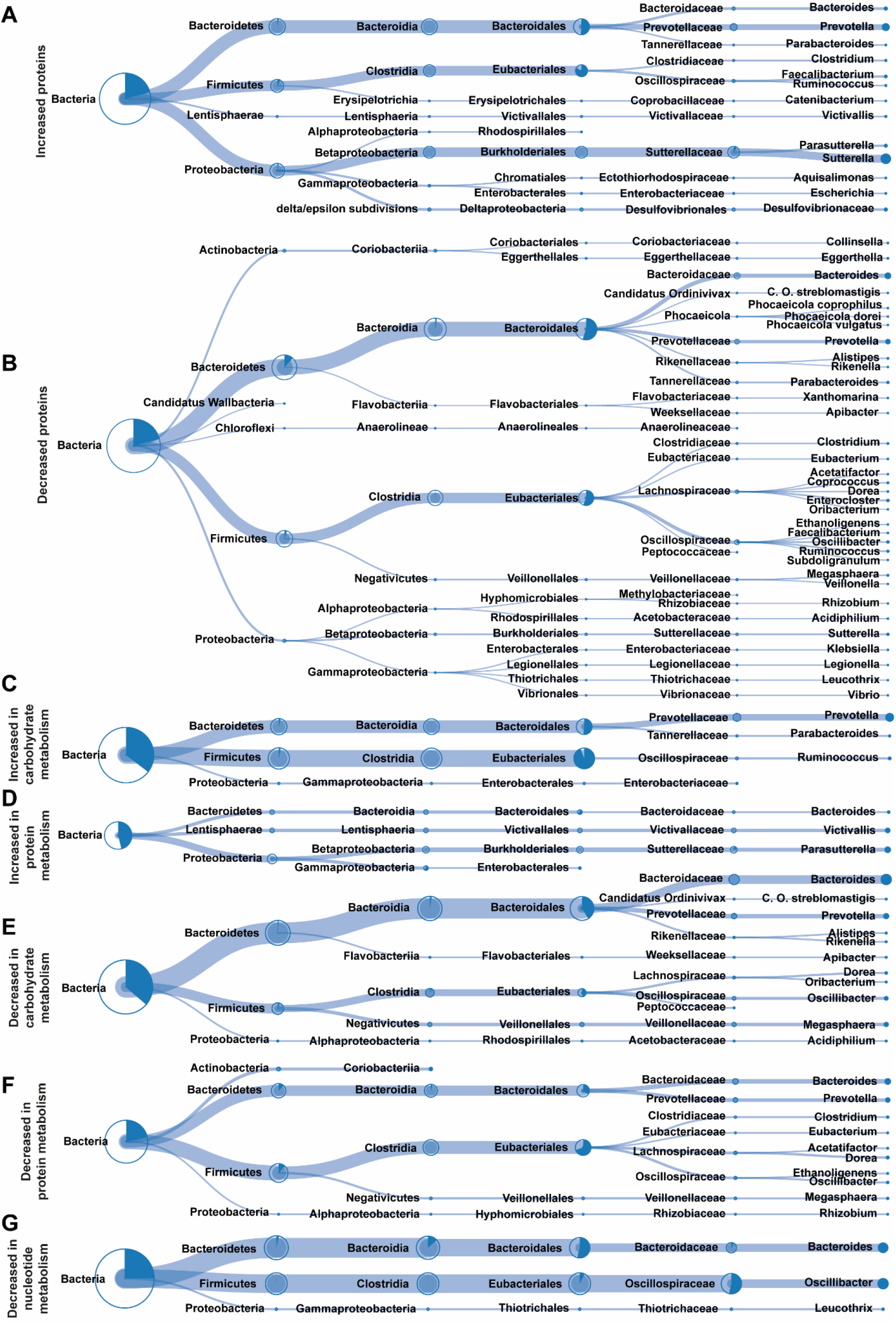
Taxonomic annotation of peptides associated with 6-mercaptopurine (6-MP) exposure. Bacterial taxonomic annotation of proteins that were increased (A) or decreased (B) in all donors due to 6-MP exposure. Bacterial taxonomic annotation of proteins that were increased (C, D) or decreased (E, F, G) due to 6-MP exposure when and related to carbohydrate metabolism (C, E), protein synthesis (D, F) or nucleotide metabolism (G). Bacterial taxa are represented up to genus level and the integrated pie charts and line thickness indicate the fraction of the respective lower taxa that could be annotated. Figures were extracted and adapted from UniPept 4.0^32^.

Increased proteins related to carbohydrate metabolism were mainly derived from Firmicutes, mainly Eubacteriales, and to a lesser extent from Bacteroidetes, especially *Prevotella*, and a few from Proteobacteria (Fig. 6C), whereas proteins related to carbohydrate metabolism that were decreased due to 6-MP exposure mainly derived from Bacteroidetes, mainly *Bacteroides* and lesser *Prevotella*, and to a lesser extent from Firmicutes, mainly Eubacteriales (Fig 6E). Increased proteins related to protein synthesis, mostly ribosomal proteins, mainly belonged to Proteobacteria, mainly *Parasutterella* (Fig. 6D), while decreased proteins mainly belonged to Firmicutes, mainly Eubacteriales, and to a lesser extent to Bacteroidetes, of which *Bacteroides* slightly dominates *Prevotella* (Fig. 6F). Finally, four of the decreased proteins are involved in nucleotide metabolism in Bacteroidetes, mainly *Bacteroides*, and Firmicutes, mainly Oscillospiraceae, and to a lesser amount in *Leucothrix*, belonging to the phylum Proteobacteria (Fig. 6G).

## Discussion

Recent studies showed that numerous medical drugs can influence growth of various intestinal bacterial strains^9^. In CD, drug-induced changes in microbiota composition and function may contribute to the reduction of mucosal inflammation or impact drug response^10^. Therefore, this study aimed to explore the effect of three CD drugs, namely budesonide, 6-mercaptopurine (6-MP), and tofacitinib, on CD patient microbiota composition and metabolism using an *ex vivo* fecal culture system.

In the present study, we found no significant effect of the selected CD drugs on the overall microbial community structure. However, when exploring the effects of these drugs on the fecal metaproteome for a single patient, profound effects were found after both, 6-mercaptopurine and tofacitinib exposure. Furthermore, extending the analyses of 6-mercaptopurine exposure to fecal samples of all five CD patients led to over- and underrepresentation of specific proteins in all donors.

In previous years, numerous studies have investigated the intestinal microbiota composition based on 16S rRNA marker gene sequencing. Thereby, an altered microbiota composition was found to be associated with CD onset and disease course as well as with treatment response^7,35,36^. When CD drugs are able to beneficially alter the microbiota composition resembling a “healthier” microbiota, this might add to the treatment response of these drugs in addition to their known direct immunomodulatory properties. For this study, we chose budesonide, 6-MP, and tofacitinib, because the former two are frequently used for CD treatment and have different working mechanisms, while tofacitinib is a new oral drug that has been found effective for CD treatment^3,18^. For all drugs, we chose (supra-)physiological concentrations that may be found in the terminal ileum and colon.

Based on previous cross-sectional clinical or *in vitro* strain culture studies, we expected to detect compositional changes induced by budesonide and 6-MP^9,12,14^, while tofacitinib has, to our knowledge, not yet been investigated. Interestingly, we could not detect any significant alterations in microbial richness, diversity and community structure. Only the *Colidextribacter* genus was significantly less abundant due to tofacitinib exposure in the higher dosage. A potential explanation for the almost complete absence of compositional changes may include that the microbial community within a fecal sample is more resilient to disturbances by xenobiotics, such as medical drugs, when compared to single strains^37^, at least for a short period of exposure. In addition, the large inter-individual variations in microbiota composition and the small number of samples, might also limit the identification of consistent shifts across individuals in response to drug exposure. Moreover, the culture period of 24 hours might be too short to detect shifts in the microbiota composition, especially since free DNA and DNA from non-viable bacterial cells will also be picked up during the analysis^22^.

Based on previous studies, we initialy expected alterations in the microbiota composition as response to the examined drugs. Together, two studies reported growth inhibition due to 6-MP of several bacterial strains, including *Prevotella corpi, Campylobacter concisus*, and *Bacteriodes fragilis*^9,12^. The explore general bactericidual effects, future studies should include absolute bacterial abundances in response to IBD drugs and confirm potential findings in more complex *ex vivo* or clinical studies.

Recently, research corroborates that the microbiota composition itself may not be the dominant driver of microbiome-host interactions. Instead, functional read-outs, such as the microbial proteome, may indicate more directly the impact of microbial changes on host physiology^8^. Therefore, for the first time, we analyzed the bacterial metaproteome in response to our selected CD drugs. When first screening each drug in one patient-derived fecal culture, the metaproteome could discriminate 6-MP and tofacitinib, but not budesonide, from the control condition. Although, 6-MP and tofacitinib did not significantly change the microbiota composition, the metaproteomic analyses clearly showed that both have a significant impact on fecal bacterial function.

Since 6-MP appeared to have the largest impact on the metaproteome, we chose to explore its impact in all five CD patient-derived fecal cultures. Intriguingly, besides an individual microbiota composition, which has often been described in previous research^7,38^, the bacterial metaproteome also appeared to be highly individual. This finding is in line with previous metaproteomic research, including other drug compounds^13^. Despite the highly individual metaproteomes, 6-MP did lead to significant and consistent over- and underrepresentation of protein clusters in all donors. The largest shifts have been annotated to bacterial carbohydrate metabolism. The affected taxa may either have de- or increased their carbohydrate metabolism in general or altered related pathways. For instance, mannose PTS system EIIAB component is increased due to 6-MP exposure. It phosphorylates D-Mannose, which could indicate an increased production of D-Mannose-6P. Since the other enzymes, which can conduct the same function (*i*.*e*. hexokinase and mannokinase) were not found to be decreased, we can assume that D-Mannose phosphorylation may indeed be increased due to 6-MP exposure. From all carbohydrate metabolism related proteins, the glycolysis/gluconeogenesis pathway seemed to be most affected. Related to this pathway, phosphoenolpyruvate carboxykinase has been annotated to different identified bacterial proteins and was decreased after 6-MP exposure. This enzyme catalyzes Oxaloacetate into Phosphoenolpyruvate, which can either be converted into Pyruvate for energy production, or into Glycerate-2P for glucose production and storage. In the same pathway, 2-oxoglutarate/2-oxoacid ferredoxin oxidoreductase subunit alpha was decreased after 6-MP exposure, which can convert Pyruvate into Acetyl-CoA, that is a substrate for short chain fatty acid (SCFA) production, including propionate and butyrate^31^. A decrease in bacterial Acetyl-CoA production may lead to a decrease in SCFA production. In the present study, downregulation of 2-oxoglutarate/2-oxoacid ferredoxin oxidoreductase subunit alpha was specifically annotated to the order of Bacteroidales, including *Bacteroides fragilis*, which produces propionate^39^. In contrast, butyrate-producing Firmicutes were not downregulated in this pathway. Consequently, 6-MP intake may increase the relative concentration of luminal butyrate, which can have an effect on the patient’s intestinal physiology and immune response^39^. However, to investigate whether the downregulation of 2-oxoglutarate/2-oxoacid ferredoxin oxidoreductase subunit alpha and phosphoenolpyruvate carboxykinase leads to less propionate production, SCFA concentrations need to be measured in the *ex vivo* cultures. In case these measurements can confirm a shift in SCFA composition due to 6-MP, SCFA composition needs to be examined in patients before and during 6-MP therapy.

Another group of proteins that was altered due to 6-MP expression includes ribosomal proteins and other proteins related to bacterial protein synthesis. Several different proteins were decreased, mainly in Firmicutes and Bacteroidetes, while other proteins were increased, mainly in Proteobacteria. Based on these results, we can assume that RNA translation may be generally decreased in Bacteroides and Firmicutes, but increased in Proteobacteria.

In general, protein expression is altered differently among bacterial taxa. Consequently, this may lead to taxon-dependent alterations in metabolite or protein production and excretion. Further research should elucidate the subsequent effects on interactions with the host.

Bacterial metaproteomics of commensal ecosystems is an innovative research field, which harbors promising and potentially clinically relevant outcomes. However, microbial proteome databases are yet mainly focused on pathogens and lack many protein and pathway annotations. For this study, only less than two third of the significantly increased or decreased proteins could be annotated using KEGG orthology. Therefore, additional interesting outcomes may have been missed. Further, the pathway annotation and subsequent interpretation needs to be taken with precaution, since the GhostKOALA tool also annotates human pathways to the microbial proteins, even though the query was specified for prokaryotes. Furthermore, taxon annotation using UniPept provides valuable insights, but is yet also limited. It creates tryptic peptides from the submitted proteins, which in turn are annotated proteins based on the peptides that were identified and analyzed by Proteome Discoverer. This additional step of data processing might introduce deviations from the original proteins and should be avoided in future, improved analysis tools. With regard to the quality of the annotated taxa, UniPept annotated some proteins to pathogens, such as *Vibrio* and *Legionella*, as well as to the marine species *Leucothrix*. These genera are very unlikely to be present in fecal samples of the study populations and were not picked up by our metagenomic analysis. Therefore, these annotations should be interpreted with precaution and rather demonstrate an interesting future analysis as soon as its reliability has been improved by at least adhering to the taxa of the submitted habitat.

Although the opportunities for microbial metaproteomic analyses need to be optimized, our study clearly demonstrates the added value of microbial metaproteomic when compared to metagenomics by unraveling functional microbial responses to common drugs. Furthermore, the rapid workflow and high-throughput expansion capacities offer promising possibilities to apply this tool in clinical drug screening protocols. Thereby, treatment choice may be co-facilitated taking into account microbial drug metabolism and subsequent harmful or complementary effects on intestinal inflammation.

To further increase the knowledge on the effect of drugs on microbiome-host interactions in intestinal inflammation, larger studies are needed which allow the analysis of individual patients and patient subgroups. The combination with clinical multi-omics studies may further pave the way for clinical translation and implementation^10^.

In conclusion, although CD drugs may not significantly change the intestinal microbial composition, they are able to significantly alter microbial protein expression within 24 hours. Thereby, the communication with the host may be altered with subsequent beneficial or harmful effects on intestinal healing. The clinical relevance of these alterations needs to be further investigated.

## Acknowledgements

HB received funding from the NUTRIM graduate program grant of Maastricht University.

